# Metric Validation for Detection of Delayed and Directed Coupling

**DOI:** 10.1101/2025.04.02.646886

**Authors:** Kate Dembny, Hafsa Farooqi, Alexander B. Herman, Theoden I. Netoff, David P. Darrow

## Abstract

**Background:** The brain functions as a complex network of billions of interconnected neurons, coordinating processes from basic reflexes to high-level cognition. Dysfunction in these networks contribute to neurological and psychiatric disorders, including epilepsy, depression, and Parkinson’s disease. Understanding these network alterations is essential for developing effective therapies. However, reconstructing network topology from human electrophysiology data is challenging due to sparse sampling, measurement noise, and variable time delays in interregional communication. Effective connectivity (EC) metrics have been developed to infer directed neural interactions, but their accuracy and robustness under real-world data constraints remain unclear. This study empirically evaluates common EC metrics to determine which most accurately reconstruct network topology in the presence of data limitations.

**Methods:** We generated Erdős–Rényi networks and simulated time series using a time-delayed vector autoregressive (VAR) model. We systematically varied network size, data length, measurement noise, and network coverage. Variations of four commonly used EC metrics, cross-correlation, Granger causality, mutual information, and transfer entropy, were evaluated for reconstruction accuracy using cosine distance to compare estimated and true connectivity matrices.

**Results:** Multivariate transfer entropy demonstrated the highest accuracy across various conditions but required significantly longer computation times. For small networks (<30 nodes), mutual information and Granger causality rapidly and accurately reconstructed networks. For larger networks, partial cross-correlation performed well with good computational efficiency. Notably, zero-lag metrics perform no better than chance in nearly all conditions.

**Conclusion:** The choice of an EC metric should consider specific data constraints.While multivariate transfer entropy is the most reliable across conditions, its long runtime limits its practical application. For large networks, partial cross-correlation offers a faster and reasonably accurate alternative. Granger causality and mutual information are effective for small networks. Critically, time-lagged metrics are essential for accurate network reconstructions, as failing to account for time delays leads to reconstructions no more accurate than random network models.

## INTRODUCTION

Neuroscience increasingly recognizes that cognitive functions arise from interactions between distributed brain regions rather than isolated activity in single areas (Beaty et al., 2016; Bressler & Menon, 2010; Bullmore & Sporns, 2009; Park & Friston, 2013; van den Heuvel & Sporns, 2013).This shift in understanding is supported by studies identifying network-level dysfunction in various neurological and psychiatric disorders, such as altered frontostriatal circuit connectivity in schizophrenia that correlates with positive symptom severity (Fornito et al., 2013; Gonzalez-Burgos et al., 2011) and increased connectivity between default mode and salience networks in individuals with depression (Mulders et al., 2015). Similarly, conditions such as Parkinson’s disease (Wu et al., 2011) and traumatic brain injury (Sharp et al., 2014), historically conceptualized as localized dysfunctions, are now understood to involve widespread network disruptions. These findings underscore the importance of inter-regional communication in understanding a broad variety of neurological and psychiatric disorders (Fornito et al., 2015; Palop et al., 2006; Vinogradov & Herman, 2015). Fundamental to understanding network connectivity is the availability of metrics that measure communication and robustly and reliably identify these network connections. In practical application, the ability to identify these connections is complicated by the size of these networks, limited and noisy data, sparse sampling of brain areas, and physiological time lags in communication.

Determining the interactions between brain regions can be approached through causal manipulations, but these methods are often impractical or unethical in human studies. Instead, we rely heavily on observational data and a wide range of analytical methods to infer communication between regions, including anatomical, functional, and effective connectivity (EC). Anatomical connectivity maps the structural pathways along which communication may occur but does not capture the dynamic nature of brain activity (Lazar, 2010). As a static model, anatomical connectivity cannot estimate dynamical changes in communication that occur over time, which are central to complex brain function (Sporns et al., 2005). Functional connectivity, long a focus of functional MRI, can measure these dynamical relationships using statistical measures (Friston, 2011) but fails to identify the directional influence necessary to understand causal relationships in complex brain networks. Effective connectivity aims to infer directed interactions while accounting for physiological communication lags (Friston, 2011; Lang et al., 2012). These communication lags provide information about the directionality of connection based on the premise that causes precede their effects in time (Valdes-Sosa et al., 2011). Consequently, EC metrics provide a powerful framework for understanding network dynamics, yet their accuracy depends on the methods used for estimation.

Numerous effective connectivity metrics have been proposed, but each comes with inherent limitations that complicate interpretation. Despite their widespread use, no consensus has emerged on which metric most accurately reconstructs neural networks across varying conditions. A key challenge is that many studies apply EC metrics without validating their performance against known ground-truth networks. Additionally, real-world electrophysiology data often contain factors that can distort connectivity estimates, including sparse sampling, measurement noise, and limited data length. To address these gaps, we systematically evaluate commonly used EC metrics by testing their ability to reconstruct simulated networks under conditions that mimic typical electrophysiological constraints. These experiments allow us to identify how well each metric performs and identify optimal conditions for use. By comparing how well each metric performs across different scenarios, we aim to identify optimal methods for reconstructing brain networks and improving the reliability of connectivity-based neuroscience research.

## METHODS

Consider the brain network with regions identified as nodes 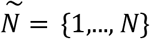, and the behavior of each node, represented by a Gaussian stationary time series, *X*= {*x*_1_,…,*x*_*N*_*}*. For the electrophysiological signal associated with each region, we discretize time with the sampling time *T*_*s*_ into discrete time samples *k*.

Therefore, the actual time *t* at each sampling instant is equal to *t* = *kT*_*s*_. Since signals are defined for finite time, we represent the final time sample with *k*_*f*_, such that *k* = *0*, …, *k*_*f*_ *− 1*.

### A. Network Simulations

In this study, we simulate networks with time-lagged communication between nodes. Each simulated network is represented by a connectivity matrix *C[m, n]* of dimensions *N ×N*, where the first dimension, *m*, of the matrix represents the source node sending information, and the second, *n*, represents the node receiving that information. A random number generator is used to create a coupling matrix with a 10% coupling probability between any two nodes, resulting in sparse and generally stable networks with Erdős–Rényi topologies.

Given this topology, we simulated time series data using coupled oscillators with time delays representing the interregional communication. Each node’s activity was modeled as a second-order coupled oscillator, incorporating lags to reflect realistic transmission delays between brain regions. The system is governed by the following equation:

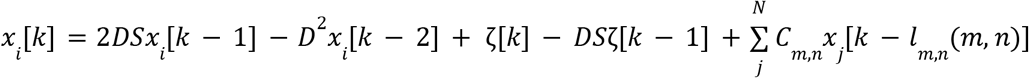

Where 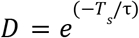 is a decay term with time constant *τ*, and *S* = *cos(ω*_*0*_ *k * T*_*s*_) determines the natural frequency of oscillation with the center frequency at *ω*_*0*_ . *ζ(t)* is a Gaussian distributed random number generator, and *l(m, n)* is the lag between nodes *m* and *n*, which is a randomly selected integer between 1 and 10 timesteps. An exemplar oscillatory system is shown in Figure 1 with the known connectivity matrix, simulated time series, and several EC reconstructions using several metrics described below.

**Fig. 1.**
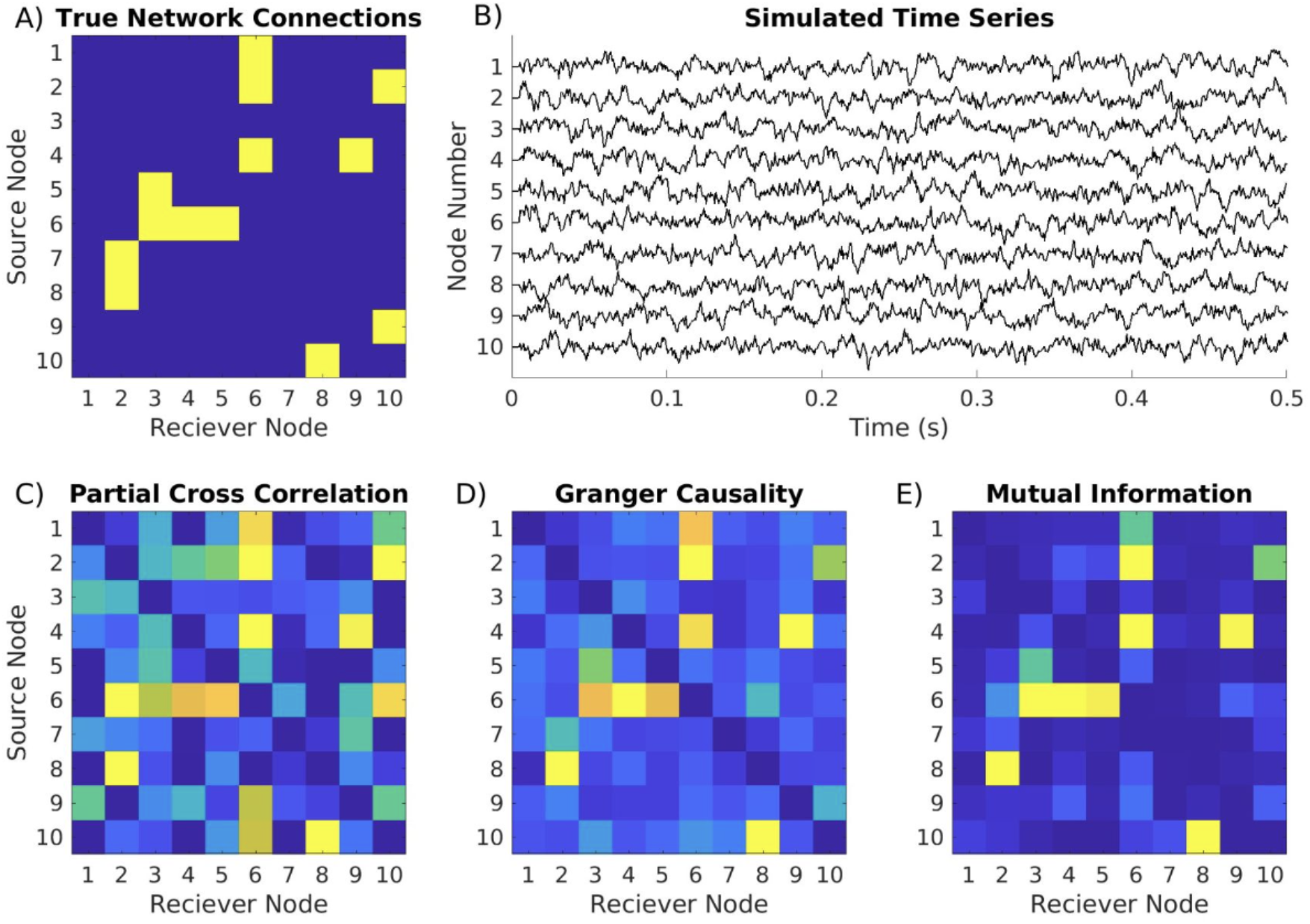
Example of oscillatory networks and reconstructions with common effective connectivity metrics. A) Adjacency matrix showing randomly generated network connectivity model, B) simulated time series generated from network in (A) containing theta frequency oscillations, C) calculated effective connectivity on time series from (B) using partial cross-correlation D) Granger causality, and E) mutual information.

While coupled oscillator models provide a biologically plausible framework for simulating network dynamics, they require predefined center frequencies and time delays, which can introduce additional complexity. Additionally, the presence of autocorrelations in these systems increases the variance in cross-correlation estimates, necessitating longer time series for statistically robust results (Jenkins & Watts, 1969). To circumvent these challenges, we implemented a vector autoregressive (VAR) model without autocorrelations. In this model, each node’s activity is driven by random noise, with short communication lags ranging from 1 to 10 time steps. The coupling strength between nodes was held constant to ensure consistency across simulations.

To establish this model, we assume that any time series signal *x*_*m*_ ∈ *X*has an equivalent *p*^*th*^ order VAR model of the form:

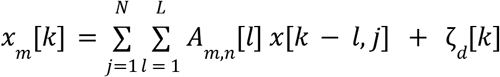

where *A* is a three-dimensional matrix of coefficients representing the coupling between node *m* and *n* at lag *l. L* represents the maximum possible lags between the nodes, and *ζ* is the dynamical noise determined by a normally distributed source with standard deviation *σ*_*d*_. The self-coupling is zero at all lags *A*_*m*,*m*_ *[1: L]* = *0*. The coupling drive from node *m* onto node *n* is a three-point filter with the coefficients [1, -2, 1] by lag, which comes from the second temporal derivative of the activity of node *j*. This approach ensures stable network dynamics while maintaining a realistic representation of time-lagged interactions.

To simulate measurement noise, we introduced an independent noise component to each node’s signal:

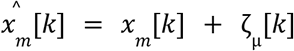

Where *x*_*m*_ is the state of the system at node *m*, and 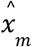 is the measured signal, including measurement noise, denoted by *ζ*_*µ*_, a normal distribution with mean zero and standard deviation *σ*_*µ*_. Simulations were generated in Mathworks 2023a. All code used for the project can be found at https://github.com/hermandarrowlab.

To illustrate the simulated networks, we generated an example 10-node network and visualized its connectivity structure. The connectivity matrix (Fig. 2A) represents the directed interactions between nodes, with nonzero entries indicating the presence of a connection. A topological representation of this network is shown in Fig. 2B, where arrows denote the direction of information flow between nodes. Time series data were generated for each node using the VAR model, incorporating predefined coupling delays and measurement noise (Fig. 2C). In these simulations, coupling strength was set at a level where inter-node communication was not visually apparent in the raw time series, ensuring that connectivity could not be inferred through simple visual inspection. Instead, effective connectivity metrics were required to reconstruct the underlying network structure, allowing for an objective evaluation of their performance.

**Fig. 2.**
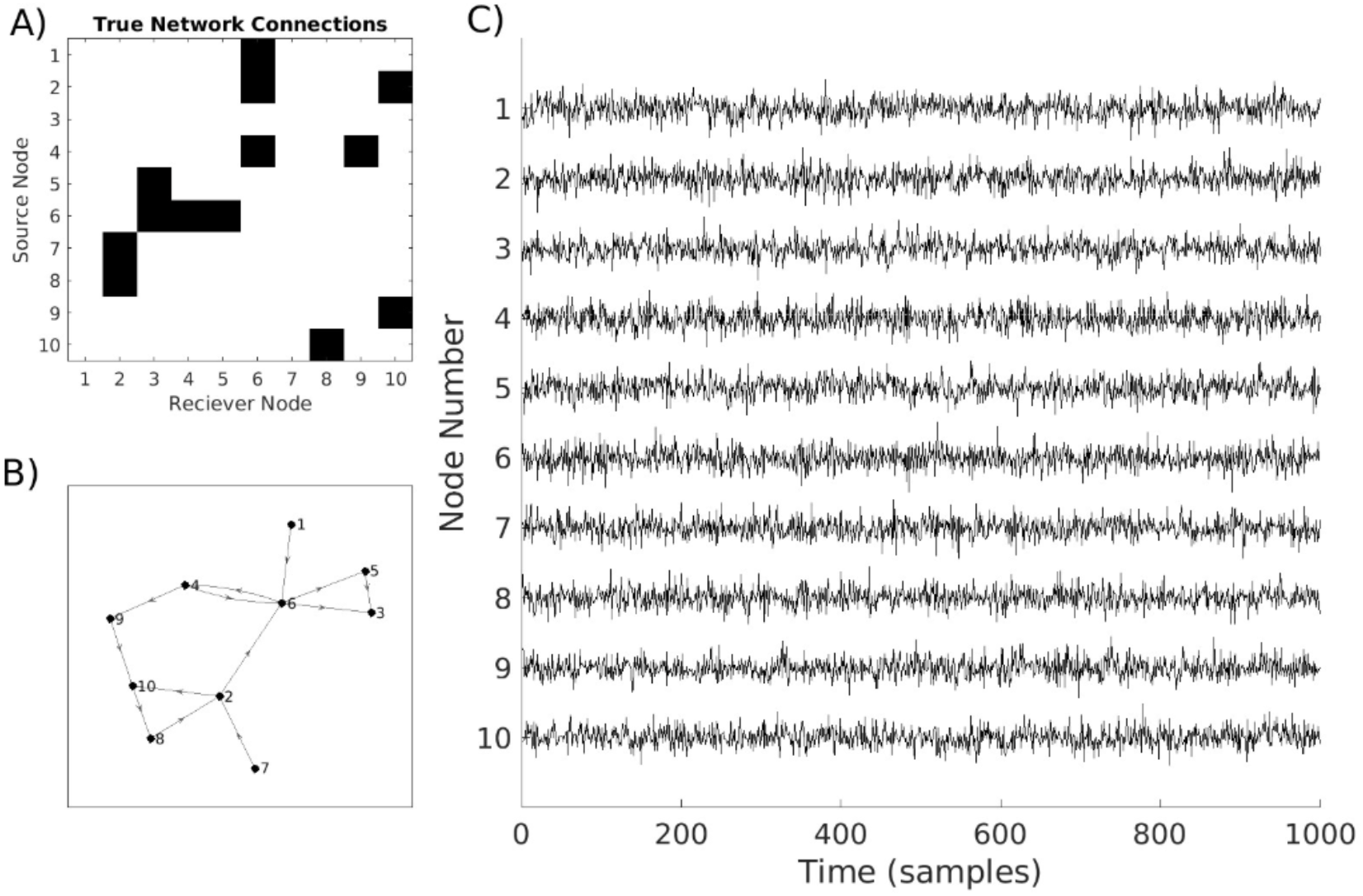
Sample of network and time series data generated by model. A) Randomly generated adjacency matrix of network connectivity. B) Visual representation of topology of the network in A), with directional arrows representing directed connection. C) Sample time series for 10-node network with time-lagged connections generated by VAR model using adjacency matrix in A).

**Fig. 3.**
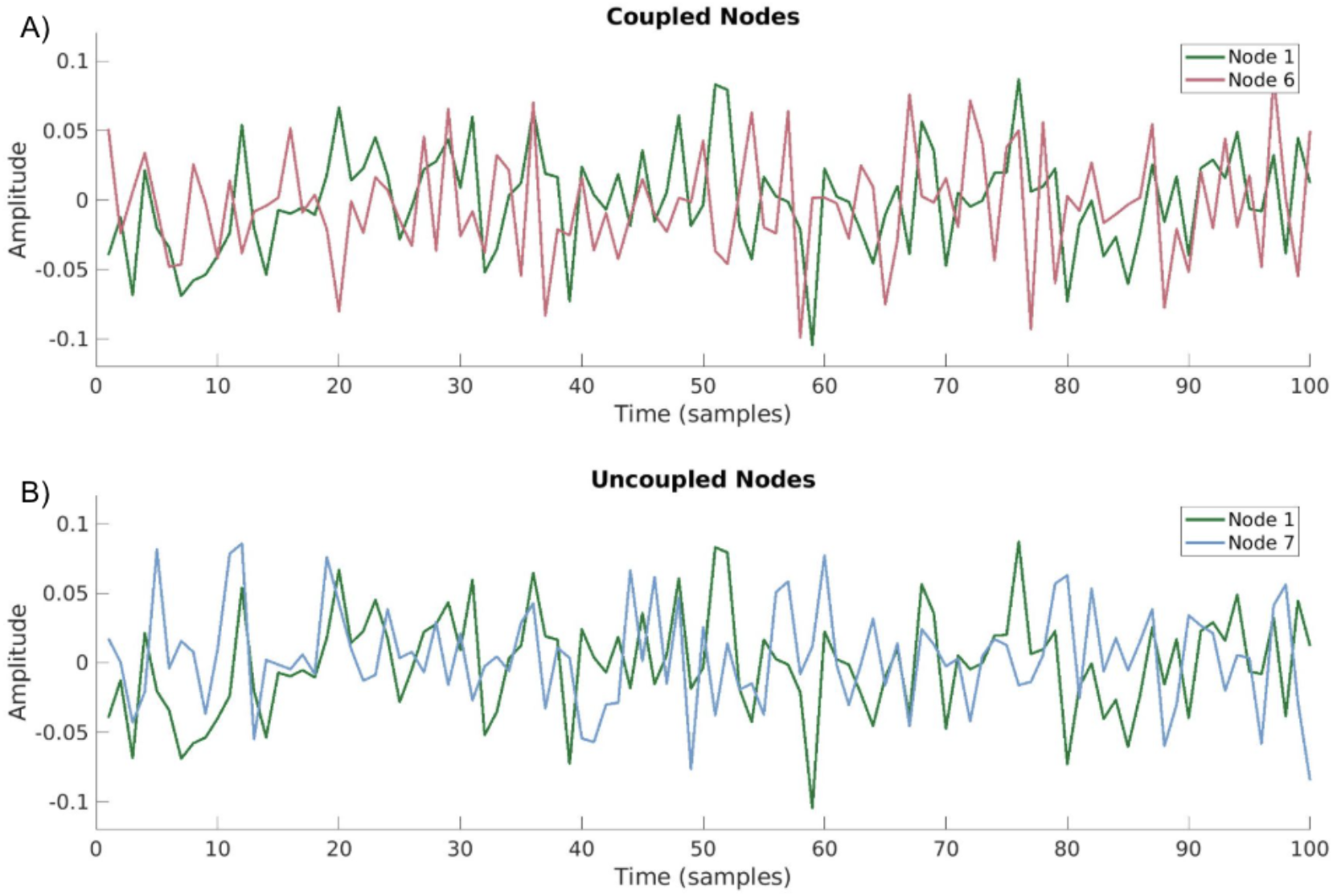
Channel coupling is not visually identifiable in coupled versus uncoupled nodes. A) Example of time series of two coupled nodes from network seen in Fig. 2. B) Example of time series of two uncoupled nodes from network seen in Fig. 2.

### B. Parameter variation

Four model parameters were varied when generating time series data (Table 1). The variations of these parameters model conditions that limit network reconstruction reliability in intracranial electrophysiology data. These parameters include:

**Table 1.** Parameter Variation. This table identifies parameter sets used to generate each set of simulated data when experimental parameters are varied.

|  |  |  |  |  |  |
| --- | --- | --- | --- | --- | --- |
| Experimental Parameter |  | Nodes | Time Points | SNR | Network Coverage |
|  | Number of Nodes | 10, 20, 30, 50, 65 | 10,000 | No Noise | 100% |
|  | Number of Time Points | 50 | 500, 1,000, 5,000, 10,000 | No Noise | 100% |
|  | SNR | 50 | 10,000 | 0.01, 0.05, 0.1, 0.5, 1, 5, 10, 50 | 100% |
|  | Network Coverage | 50 | 10,000 | No Noise | 100%, 90%, 80%, 70%, 60%, 50%, 40%, 30%, 20%, 10% |

### Number of Nodes (N)

Networks with 10, 20, 30, 50, and 65 nodes were generated. When other parameters were varied, the number of nodes used for the simulation was held constant at 50.

#### Number of Data Points (k)

The number of data points in the time series was varied with options of 500, 1,000, 5,000, and 10,000. The number of time points was held constant at 10,000 when other parameters were varied.

#### Measurement Noise (U)

Independent white measurement noise was added to each node as a function of signal-to-noise ratio (SNR) using Matlab’s awgn function. SNR values were 0.01, 0.05, 0.1, 0.5, 1, 5, 10, and 50. Simulations contained no noise when other parameters were varied.

#### Percent Network Coverage

Networks were simulated with all of the default parameters above. A pseudorandom number generator selected 5 nodes (10% of the network) and their time series to exclude from the network and EC calculations. This was done iteratively until only 5 nodes remained in the network (10% network coverage). Network coverage was held at 100% when other parameters were varied.

### C. Network Reconstruction Metrics

In this work, we evaluate the ability of standard effective connectivity (EC) metrics to identify time-lagged and directed communication from time series data. We focus on four metrics: cross-correlation, Granger causality, mutual information, and transfer entropy. All of these methods have implementations available as packages in Matlab or Python. Below, we briefly define the mathematical formulations for each metric.

Consider two measured discrete time series signals 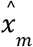 and 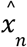, where 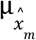 and 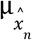 are the mean of the brain signals 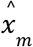 and 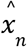, respectively, and 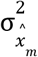 and 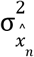 are their variances. When analysis excludes time series from 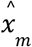 and 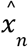, we denote the remainder network as 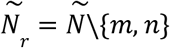 and the remainder time series as 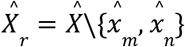.

#### Cross-Correlation

Correlation is a well-established measure that estimates the linear relationship between two signals and is robust even in noisy conditions (Netoff et al., 2004; Ostojic et al., 2009; Rodu et al., 2018). Cross-correlation can be measured between two signals at different time delays to identify communication lag. These lags provide information about communication direction given the assumption that causes precede their effects. However, there are limitations. Cross-correlation assumes linear relationships between time series, so it may miss nonlinear relationships (Sakkalis, 2011) with no significant linear component. Additionally, the metric has difficulty identifying bidirectional interactions and struggles to interpret multiple delays with high correlation values (Bastos & Schoffelen, 2016).

#### Bivariate Cross-Correlation

In the time domain, the cross-correlation of 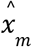 and 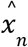 at lag *l* can be calculated using the following mathematical formulation:

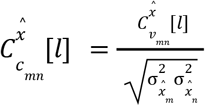

where covariance 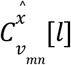 between the brain signals 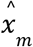 and 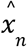 is further defined as:

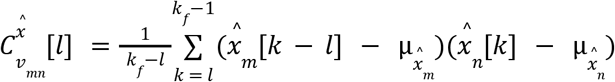

The cross-correlation is equivalent to normalized cross-covariance.

#### Partial Cross-Correlation

Within a time series, it is important to identify whether similar recorded signals result from communication between two connected regions or, alternatively, if the two signals have a shared source with which they both independently communicate. In an attempt to resolve this conflict, partial cross-correlation has been proposed.

When calculating the partial lagged cross-correlation in the time domain, information contained in 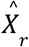, can be used to estimate the best linear predictor 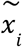 of the signal 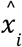 that minimizes the residual 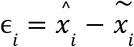. The partial lagged cross-correlation of the time series signals 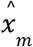 and 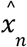 is defined as the lagged cross-correlation between the residuals *ϵ*_*m*_ and *ϵ*_*n*_.

In practice, however, it is difficult to identify accurate estimators of 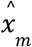 and 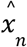 in the time domain. We instead calculate cross-correlation in the frequency domain, where the need to calculate these estimators is avoided. To accomplish this frequency domain calculation, for any frequency *λ* ∈ *[− π, π]*, we define the Discrete-Time Fourier transform of the covariance between the time series 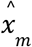 and 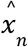as:

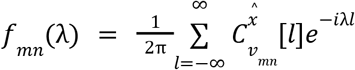

Next, the partial cross-spectral density between the signals is defined as:

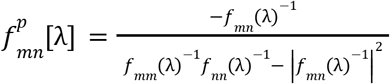

and the partial covariance 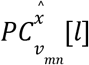 is calculated using the inverse Discrete Fourier Transform:

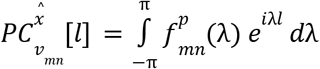

In summary, 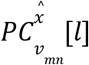 can be calculated from cross-spectral densities of the simple lagged cross-correlations *fmn (λ)* between 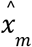 and 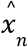 (Eichler et al., 2003; Salvador et al., 2005). Cross-correlation was calculated using the Functional Connectivity Toolbox (Zhou et al., 2009).

#### Granger Causality

Granger causality (GC) estimates the reduction in forecasting error resulting from including the history of an additional node in an autoregressive model used to estimate the next time point (Granger, 1969; Lütkepohl, 2005). Initially proposed for economics data, the metric is broadly utilized within neuroscience literature (Barnett & Seth, 2014; Shojaie & Fox, 2022). However, GC assumes the recorded time series offers a complete representation of the system, which is often untrue in neurophysiology recordings (Shojaie & Fox, 2022). While the original GC metric assumes linear relationships, nonlinear and nonparametric extensions have been developed (Barnett & Seth, 2014; Dhamala et al., 2008; Sysoeva et al., 2014; Zhang et al., 2016). To estimate GC, we assume any time series signal 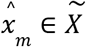 has an equivalent *p^th^* order VAR model of the form:

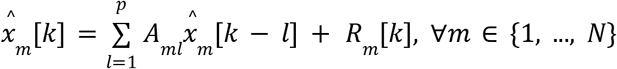

where *A*_*ml*_ are regression coefficients and *R*_*m*_ *[k]* is the residual at time sample *k*.

If the inclusion of 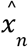 reduces the prediction error, it can be concluded that the 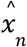 is sending directed communication to 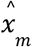. In order to calculate GC between two time series, we define the full 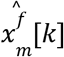 and reduced 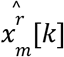 regression models of the time-series signal 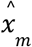:

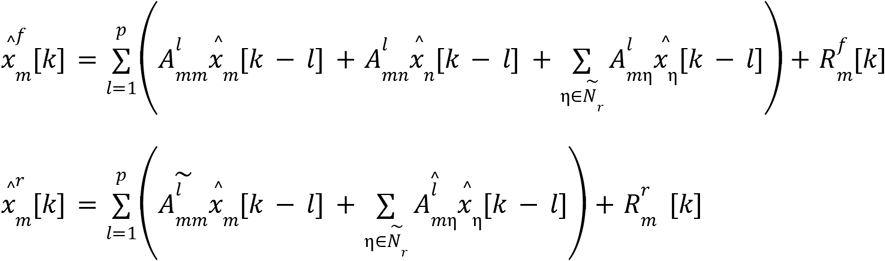

Having defined the appropriate models, we define multivariate Granger causality from 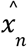 to 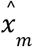 conditioned on 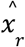 as the following log-likelihood ratio:

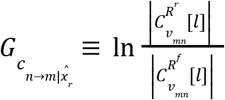

GC was calculated using the Multivariate Granger Causality Toolbox by Anil Seth (Barnett & Seth, 2014).

#### Mutual Information

Mutual Information (MI) has a mathematical basis in Shannon entropy, which estimates information flow between two signals using their probability distributions (Emmert-Streib & Dehmer, 2009; Grassberger et al., 1991). As a symmetric operator, MI calculates coupling without directionality. However, time-lagged versions have been developed by calculating MI while shifting the temporal alignment between signal processes (Schreiber, 2000), and one such implementation is used in this paper. In time series data, the assumption of independence between time samples is invalid since time series often show temporal correlations. To account for this, we assume each sampled pair *x*_*m*_ *[k], x*_*n*_ *[k]* is a random variable for every time sample. Under this assumption, we define MI as (Schreiber, 2000; Shannon, 1948):

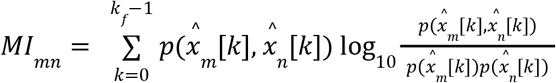

The notation 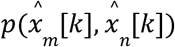 denotes the joint probability density function of 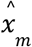 and 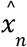, while 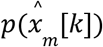 and 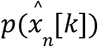 are the marginal probability density functions of 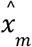 and 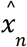 respectively at sample *k*. For lag *l*, we define the lagged MI as:

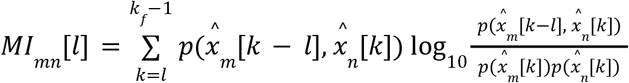

Fieldtrip was used to calculate bivariate mutual information using their `gcmi` method (Oostenveld et al., 2011).

#### Transfer Entropy

Transfer entropy (TE) utilizes Shannon entropy to quantify information transferred from one node to another (Schreiber, 2000; Staniek & Lehnertz, 2008) by calculating transition probabilities and determining whether including the history of an additional signal process improves the prediction of these probabilities. TE makes no linearity assumptions and provides directional information about interactions, though this comes at the expense of computational efficiency (Sabesan et al., 2009; Vicente et al., 2011). The absence of assumptions about the underlying relationships between nodes makes TE an attractive option for understanding biological systems that are not well characterized.

#### Bivariate Transfer Entropy

TE measures transition probabilities to evaluate directional information transfer. Consider the time series 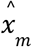, approximated by a stationary Markov process of order *l*. The conditional probability of finding 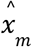 in state 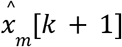 at time sample *k + 1* is calculated as

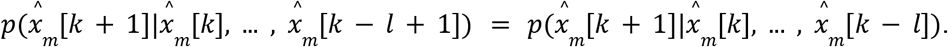

We denote 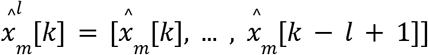. The average number of bits needed to encode one additional state of 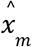, if all the previous states are known, is given by entropy rate:

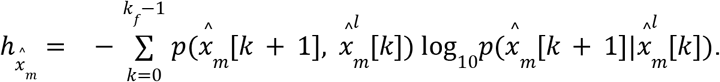

For the two time series, it is preferable to measure the deviation from the generalized Markov property 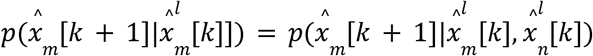 . Transfer entropy measures the influence of the states of 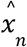 on the transition probabilities of 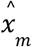, which we define:

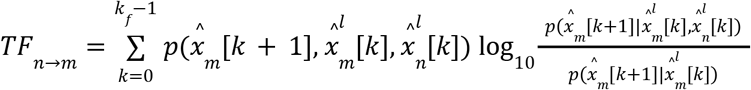

TE measures the amount of uncertainty reduced in future values of 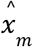 by knowing the past states of 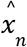.

#### Multivariate Transfer Entropy

In a multivariate setting, time series information from the residual time series 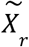 is also taken into account, and multivariate transfer entropy is calculated not only on the values of the source information 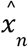 but also conditions on the information contained in the residual time series 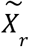. We define the time samples at *k* of the residual time series 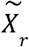 by the set 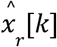 To reduce computational time, the toolbox IDTxl reduces the number of nodes included in 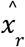, only including relevant nodes, which we define here as 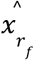. Next, multivariate transfer entropy (mTE) is defined as (Schreiber, 2000):

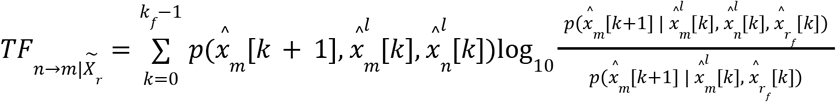

For this paper, the Information Dynamics Toolkit XL (IDTxl) toolbox was used to calculate both bivariate and multivariate transfer entropy. The `JidtGaussianCMI` estimator was used (Wollstadt et al., 2019). We note that this method can take a very long time to calculate, which was a limiting factor for our analyses.

#### Shuffled Networks

To measure statistical significance of outcomes relative to the null hypothesis that the networks extracted are random, we created random network topologies and compared them to the true networks to generate a distribution for the null hypothesis. Shuffled arrays were randomly generated using the same protocol used to generate the original networks.

### D. Runtime calculation

All runtime calculations measure the total runtime for the calculation when run on cores at the University of Minnesota’s Supercomputing Institute. These runtimes include the actual time to calculate the metrics but exclude formatting or saving data. Calculations were run on MSI’s Agate and Mangi clusters. Agate nodes have AMD 7763 processors, which run 2.5 double-precision teraflops per core at base frequency (Trader, 2021). Mangi nodes have AMD ROME processors, which run 0.043 teraflops per core at base frequency (*AMD EPYC*^*TM*^ *7002 Series Processors*, n.d.).

### E. Network similarity measure

To evaluate the accuracy of network reconstructions, we quantified the cosine distance between the estimated and true connectivity matrices. Cosine distance was chosen because EC metric outputs are arbitrarily scaled, and this measure assesses structural similarity independent of absolute values. It calculates the angle between two vectors, with smaller angles indicating greater similarity. Self-connections, represented by diagonal values in the connectivity matrices, were excluded from the analysis to prevent bias. The remaining values were flattened into a one-dimensional vector, and cosine distance was computed using MATLAB’s *pdist2* function. The cosine distance metric ranges from 0 to 2, where values closer to 0 indicate stronger agreement between the reconstructed and ground-truth networks.

### F. Statistics

Groups were compared with a Kruskal-Wallis test using Matlab’s `kruskalwallis`. If significant differences were identified, post-hoc Dunn-Sidak testing was done using Matlab’s `multcompare`. Bonferroni correction was performed to correct for multiple comparisons among groups. The number of comparisons was calculated by adding the number of metrics compared to the number of varied parameters in the model (i.e., 5 different node counts when parameter was varied + 10 EC metrics, so for statistics on node count data, = 0.05/15 = 0.003). As a result, the corrected alpha values for each condition are as follows: number of nodes = 0.003, number of time points = 0.00357, noise = 0.00277, and network coverage = 0.0025.

## RESULTS

We empirically evaluated the accuracy of several commonly used effective connectivity metrics to reconstruct simulated networks in the context of several practical limitations in intracranial electrophysiology data. An exemplar network topology of true network connectivity and its respective reconstructions can be seen in Fig. 4.

**Fig. 4.**
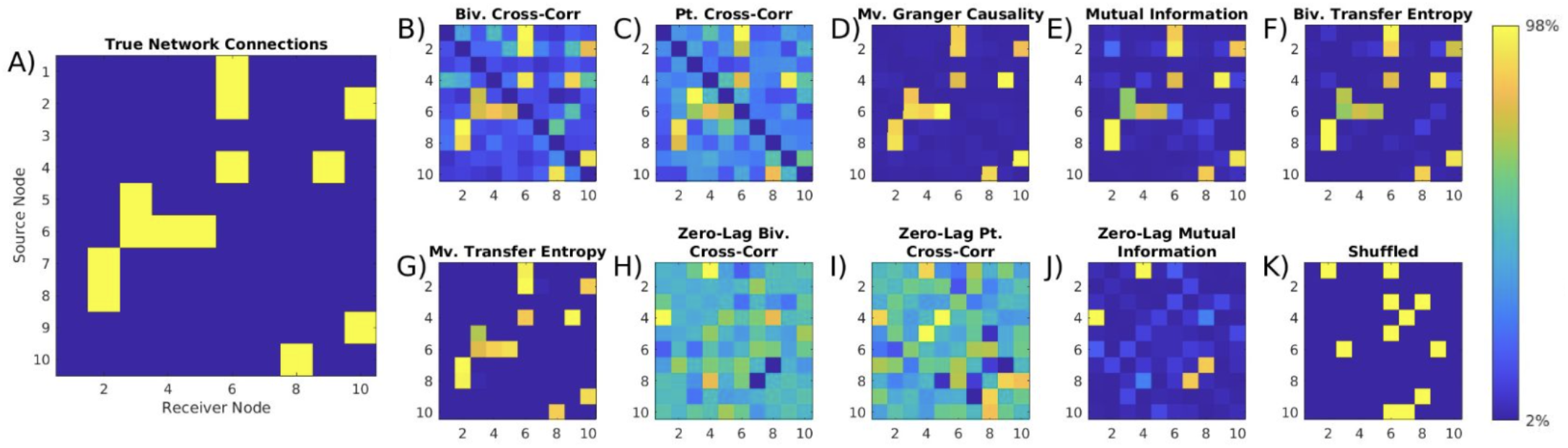
Example network reconstructions with effective connectivity metrics. Metrics are not on the same scale, so color scaling is set with the 2nd and 98th percentiles for each adjacency matrix as the maximum and minimum. A) denotes the true network connections used to generate the time series data, while the following matrices are the effective connectivity between nodes calculated using B) multi-lag bivariate cross-correlation, C) multi-lag partial cross-correlation, D) multivariate Granger causality E) multi-lag bivariate mutual information, F) bivariate transfer entropy G) multivariate transfer entropy H) zero-lag bivariate cross-correlation, I) zero-lag partial cross-correlation, J) zero-lag bivariate mutual information, K) Shuffled network with 10% coupling probability. In this figure, we observe an example network topology and metric network reconstructions of the exemplar network. We see that time-lagged metrics generate accurate reconstructions of the ten-node network, whereas zero-lag metrics visually fail to detect connections. This highlights the necessity of utilizing metrics that can successfully identify connections in systems where delays are present.

### Number of Nodes

Network size is crucial in intracranial electrophysiology, particularly as technological advances allow for greater numbers of intracranial electrodes to cover additional brain regions. To assess the impact of network size on reconstruction accuracy, we varied the number of nodes while holding the connection probability constant at 10%. The number of time points was fixed at 10,000 with no measurement noise.

Increasing the network size worsened most time-lagged metrics, except partial cross-correlation, which improved with increasing network size (Fig. 4B, p < 0.001). Granger causality showed the most significant decline in similarity as network size increased (Fig. 5C). At the same time, multivariate transfer entropy (MVTE) exhibited only a slight but significant decline (Fig. 5F). For 10-node networks, multivariate Granger causality (MVGC), bivariate transfer entropy (BVTE), and MVTE performed similarly (Supp. Fig. 1). However, as networks grew MVTE maintained strong performance, while GC and BVTE declined. As a result, multivariate transfer entropy outperformed Granger causality and bivariate transfer entropy in larger networks (p < 0.001 for BVTE and GC in 50-node networks). Partial cross-correlation, conversely, performed poorly in small networks but improved and showed comparable performance to MVTE in networks with 50+ nodes, with both significantly outperforming other metrics.

**Fig. 5.**
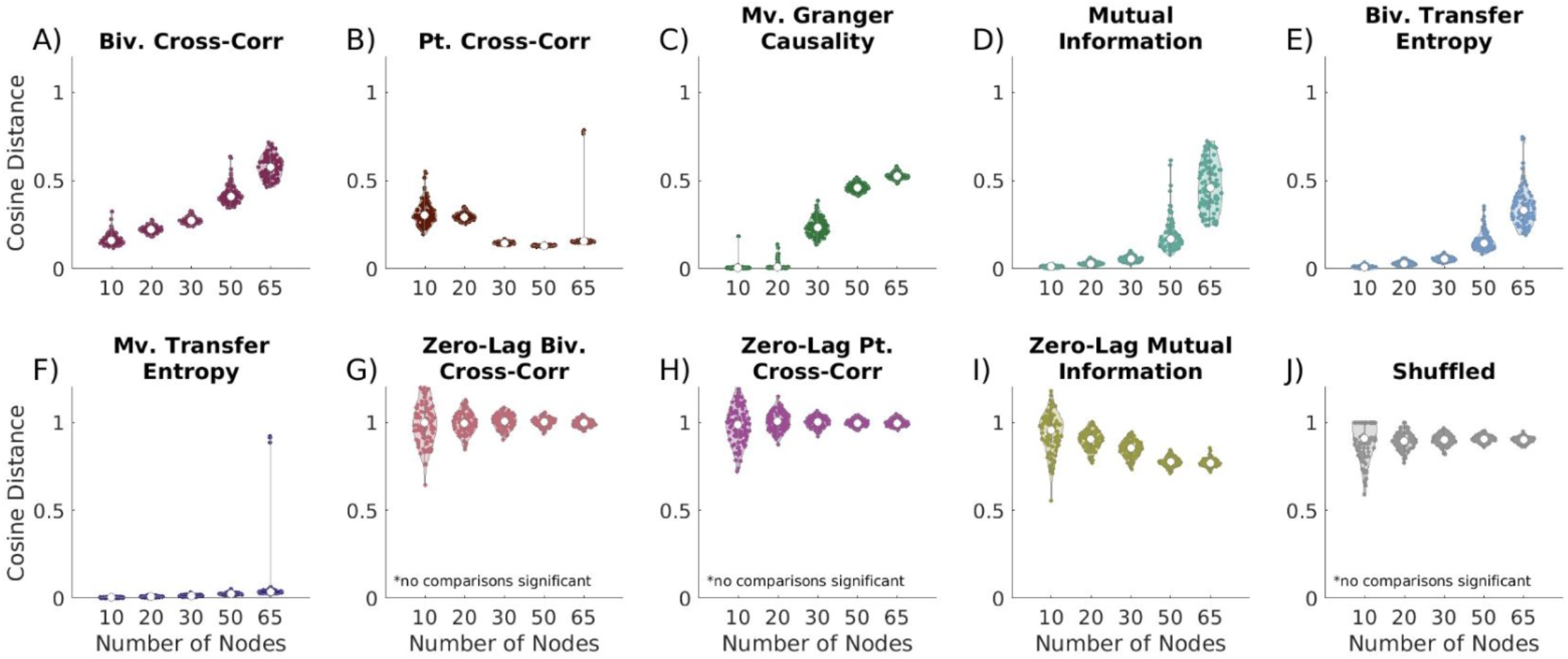
Effect of increasing network size on EC network reconstruction similarity to true networks. Cosine distance values close to 0 indicate the greatest possible similarity to the true networks. A) time-lagged bivariate cross-correlation (BVXC), all comparisons are significant; B) time-lagged partial cross-correlation (PTXC), all comparisons significant except 10 vs. 20 node networks; C) time-lagged multivariate Granger causality (MVGC), all comparisons significant excluding 10 vs. 20 node networks; D) time-lagged bivariate mutual information (BVMI), all comparisons are significant; E) time-lagged bivariate transfer entropy (BVTE), all comparisons are significant; F) time-lagged multivariate transfer entropy (MVTE), all comparisons are significant; G) zero-lag bivariate cross-correlation, no comparisons are significant; H) zero-lag partial cross-correlation, no comparisons are significant; I) zero-lag bivariate mutual information, all comparisons significant excluding 10 vs. 20 nodes and 50 vs. 65 nodes; J) shuffled, no comparisons are significant. All lagged metrics worsen with increasing network size, excluding partial cross-correlation. For 10-20 node networks, multivariate Granger causality, bivariate transfer entropy, and multivariate transfer entropy perform comparably, while outperforming other metrics. For largest network sizes, multivariate transfer entropy and partial cross-correlation significantly outperform other metrics.

Zero-lag metrics consistently underperformed or performed no better than shuffled networks across all network sizes (Fig. 5G–J). These shuffled networks serve as a negative control, representing random networks with similar structure, establishing a baseline for comparison. In contrast, time-lagged metrics consistently outperformed shuffled networks. Additionally, multivariate versions of transfer entropy and cross-correlation generally outperformed their bivariate counterparts, with the performance gap becoming more pronounced in larger networks. This suggests that incorporating additional network-wide information enhances reconstruction accuracy particularly as network complexity increases.

### Number of Time Points

Time series length is a critical factor in network reconstruction, as some studies rely on extended recordings while others analyze brief windows within behavior tasks. To assess its impact, we varied the number of time points in a fixed 50-node, measurement noise-free network. Increasing the number of time points improved all time-lagged metrics (p < 0.001 for 500 vs. 10,000 points, Fig. 6A-F), though the extent of improvement varied by metric. Most showed slight improvements, while partial cross-correlation improved significantly. Bivariate and multivariate transfer entropy performed similarly for shorter time series (500-1,000 points) (Supp. Fig. 2A-B, p = 0.067, p = 0.086). However, at 5,000 points, multivariate transfer entropy outperformed all other metrics (Supp. Fig. 2C), and at 10,000 time points, multivariate transfer entropy and partial cross-correlation performed similarly (Supp. Fig. 2D, p = 0.012). Zero-lag metrics performed poorly in all conditions, and increasing time series length did not improve the accuracy of zero-lag bivariate or partial cross-correlation, underscoring that these metrics lack information about the true networks.

**Fig. 6.**
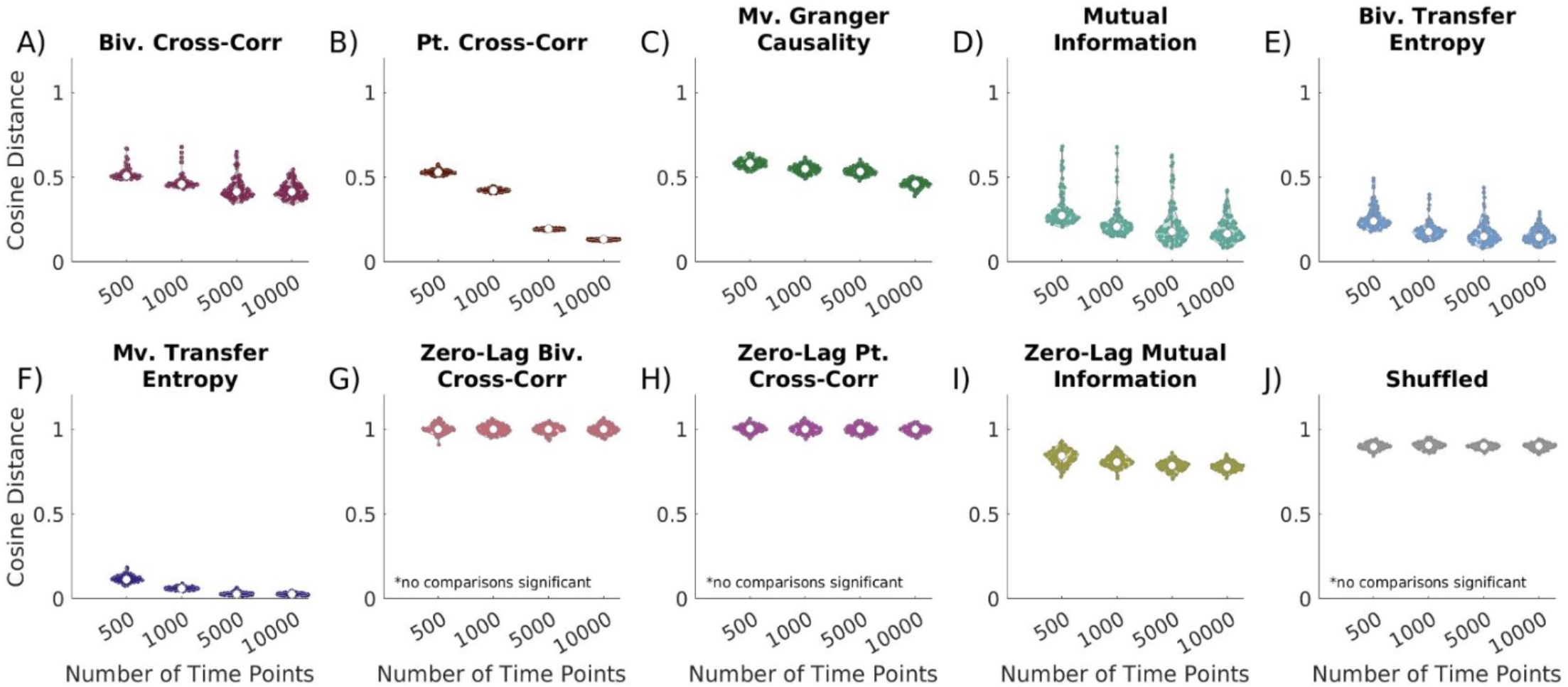
Effect of increasing time series length on EC network reconstruction similarity to true networks. Cosine distance values closest to 0 indicate the greatest similarity to the true networks. A) time-lagged bivariate cross-correlation, all comparisons significant except 5,000 vs. 10,000 time points; B) time-lagged partial cross-correlation, all comparisons significant; C) time-lagged multivariate Granger causality, all comparisons except 1,000 vs 5,000 time points are significant; D) time-lagged bivariate mutual information, comparisons of 500 vs. all other numbers of time points are significant; E) time-lagged bivariate transfer entropy, all comparisons except 1,000 vs 5,000 time points and 5,000 vs 10,000 time points are significant; F) time-lagged multivariate transfer entropy, all comparisons excluding 5,000 vs. 10,000 time points are significant; G) zero-lag bivariate cross-correlation, no comparisons significant; H) zero-lag partial cross-correlation, no comparisons significant; I) zero-lag bivariate mutual information, all comparisons except 5,000 vs. 10,000 time points are significant, J) shuffled, no comparisons significant. For all time series lengths multivariate transfer entropy is among the top performers. For 500 and 1,000 time points, bivariate and multivariate transfer entropy perform comparably. For 10,000 time point time series partial cross-correlation performs comparably to multivariate transfer entropy.

#### Measurement Noise

Noise is a common issue in neural recordings, often obscuring brain region and circuit activity while contaminating effective connectivity estimates (Bastos & Schoffelen, 2016; Pesaran et al., 2018). To evaluate its impact on network reconstructions, we introduced varying levels of measurement noise into time series data, altering the signal-to-noise ratio (SNR). Simulations were conducted with 50 nodes, 10,000 time points, and a constant 10% connection probability.

All time-lagged methods, except bivariate cross-correlation, significantly improved as SNR increased (all p’s < 0.0001). In high-noise conditions, multivariate transfer entropy performed comparably to bivariate transfer entropy but outperformed all other metrics. When 5 *≤* SNR *≤* 10, multivariate transfer entropy consistently outperforms all other metrics. However, when SNR = 50, MVTE performs comparably to bivariate transfer and bivariate mutual information. Across all noise levels, bivariate transfer entropy performed similarly to partial cross-correlation and bivariate mutual information (Supp. Fig. 3).

As in previous tests, zero-lag metrics performed poorly at all noise levels, showing no significant difference from shuffled networks (Fig. 7G-J).

**Fig. 7.**
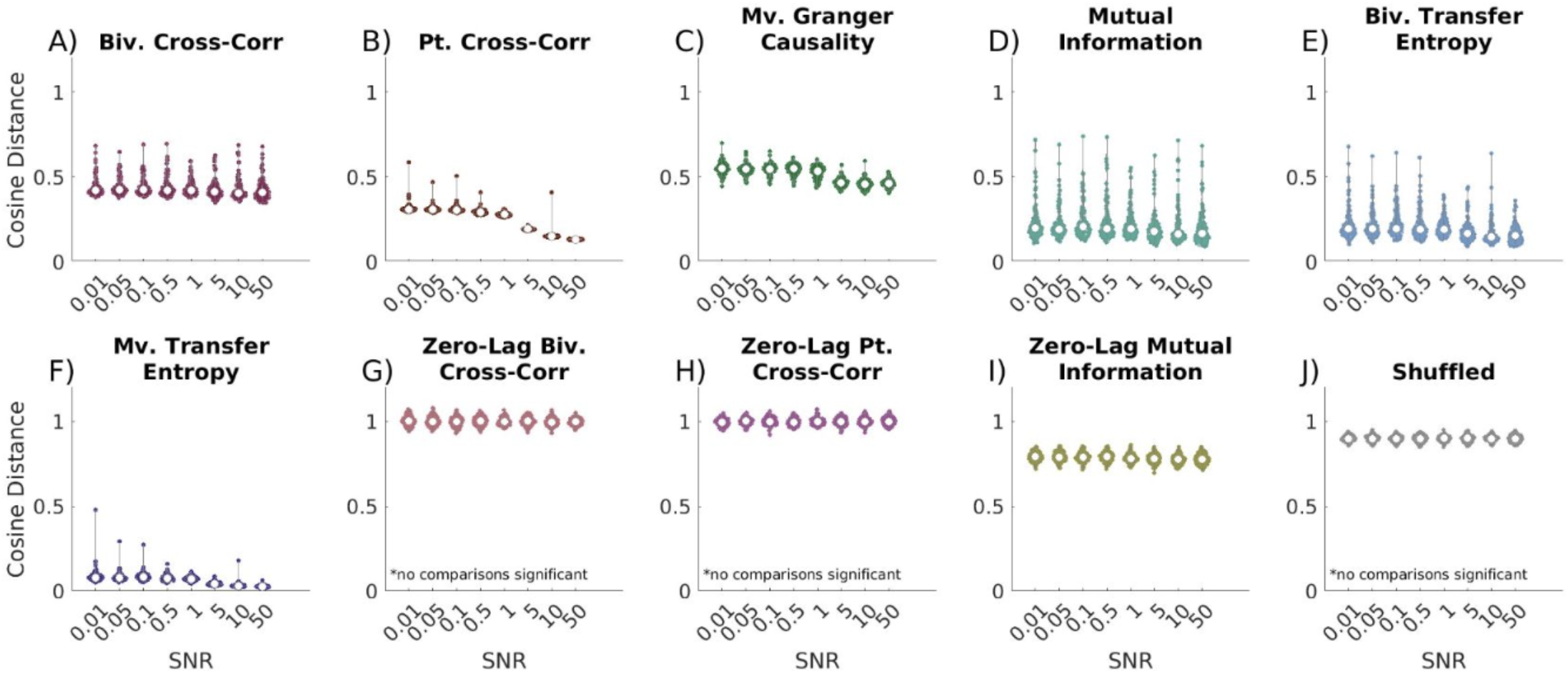
Effect of increasing SNR on EC network reconstruction similarity to true networks. Cosine distance values closest to 0 indicate the greatest similarity to true networks. A) time-lagged bivariate cross-correlation, only SNR = 0.1 vs. 10 significant; B) time-lagged partial cross-correlation, all comparisons significant except SNR = 0.01 vs 0.05, 0.01 vs. 0.1, 0.05 vs. 0.1,0.5 vs. 1,5 vs. 10, and 10 vs. 50; C) time-lagged multivariate Granger causality, significant comparisons include SNR = 0.01 vs. 5, 0.01 vs. 10, 0.01 vs. 50, 0.05 vs. 5, 0.05 vs. 10, 0.05 vs. 50, 0.1 vs. 0.5, 0.1 vs. 5, 0.1 vs. 10, 0.1 vs. 50, 0.5 vs. 5, 0.5 vs. 10, 0.5 vs. 50, 1 vs. 5, 1 vs. 10, 1 vs. 50; D) time-lagged bivariate mutual information, significant comparisons include SNR = 0.01 vs. 10, 0.01 vs. 50, 0.1 vs. 5, 0.1 vs. 10, and 0.1 vs. 50; E) time-lagged bivariate transfer entropy, significant comparisons include SNR = 0.01 vs. 5, 0.01 vs. 10, 0.01 vs. 50, 0.05 vs. 5, 0.05 vs. 10, 0.05 vs. 50, 0.1 vs. 5, 0.1 vs. 10, 0.1 vs. 50, 0.5 vs. 10, 0.5 vs. 50, 1 vs. 5, 1 vs. 10, 1 vs. 50; F) time-lagged multivariate transfer entropy, all comparisons significant except 0.01 vs. 0.05, 0.01 vs. 0.1, 0.01 vs. 0.5, 0.01 vs. 1,0.05 vs. 0.1,0.05 vs. 0.5, 0.05 vs. 1, 0.1 vs. 0.5, 0.1 vs. 1, 0.5 vs. 1,5 vs. 10, 10 vs. 50; G) zero-lag bivariate cross-correlation, no comparisons significant; H) zero-lag partial cross-correlation, no comparisons significant; I) zero-lag bivariate mutual information, significant comparisons include SNR = 0.01 vs. 10, 0.05 vs. 10, 0.1 vs.10, 0.5 vs. 10; J) shuffled, no comparisons significant. All lagged metrics and zero-lag mutual information improve with increasing SNR. For low SNR conditions, multivariate transfer entropy perform significantly outperforms all other metrics, excluding bivariate transfer entropy, which performs comparably. As noise increases, multivariate transfer entropy begins to significantly outperform bivariate transfer entropy. This is occurs when SNR = 0.1,0.5, 5, 10, and 50. Otherwise multivariate transfer entropy significantly outperforms all other metrics in all conditions. Bivariate transfer entropy performs statistically no different than partial cross-correlation and mutual information for all noise levels.

### Network Coverage

Coverage of brain networks in intracranial electrophysiology is often constrained by factors such as the limited intracranial space available for electrode placement, the cost of electrodes, the impracticality of high-density sampling, and hardware requirements for signal amplification. To assess how effective connectivity metrics performed under limited brain region coverage, we simulated partial network sampling using 50-node networks with 10,000 time points and no measurement noise. We progressively restricted the available data for reconstruction and compared the estimated connectivity to true connectivity.

Metric performance varied with declining network coverage. Bivariate cross-correlation, mutual information, and bivariate transfer entropy were unaffected (Fig. 8A,C,E), although the variance of bivariate metrics increased significantly with declining network coverage (p < 0.0001). In contrast, partial cross-correlation (Fig. 8B, p < 0.0001) and multivariate transfer entropy (Fig. 8F, p < 0.0001) exhibited significant declines in performance. Interestingly, multivariate Granger causality improved as network covered decreased (Fig. 8D, p < 0.0001), suggesting that its sensitivity to spurious connections may be reduced in more sparsely sampled networks.

**Fig. 8.**
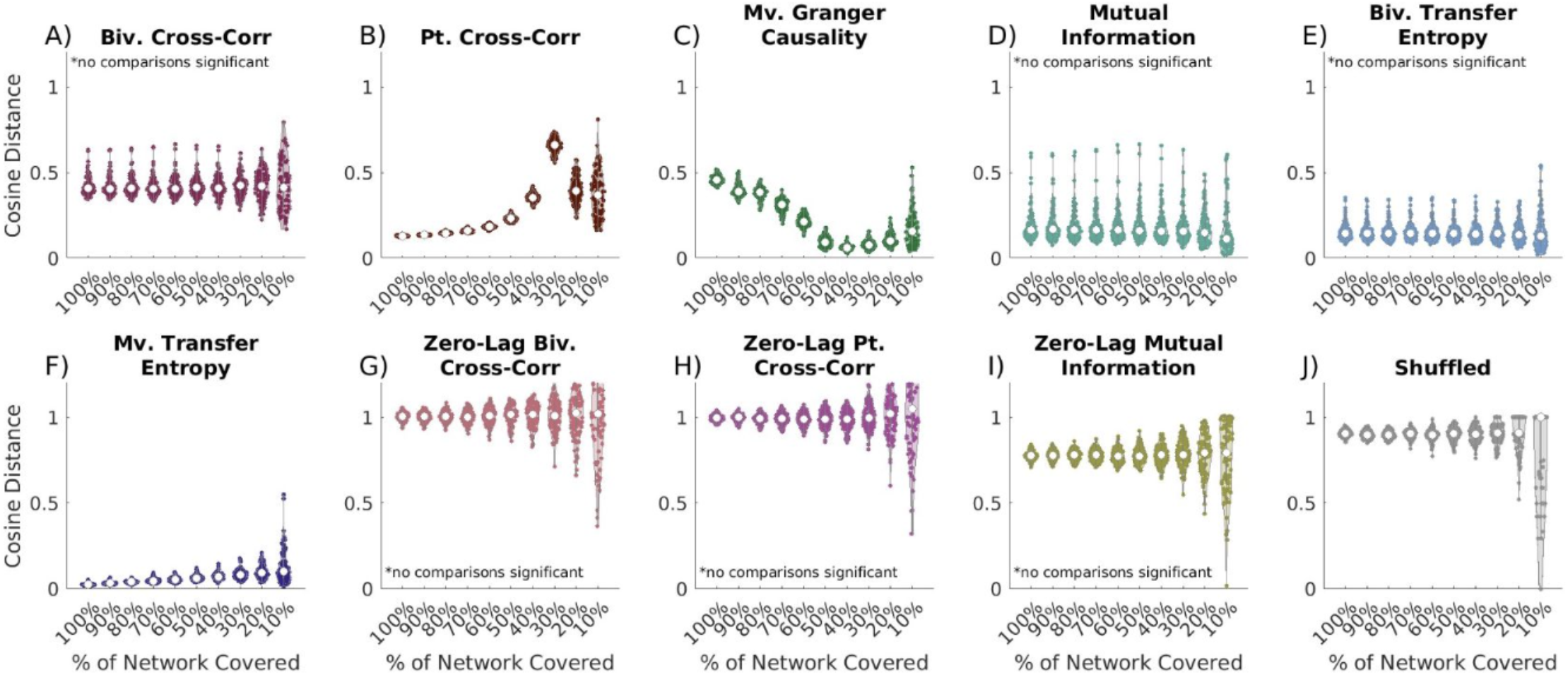
Effect of decreasing network coverage on EC network reconstruction similarity to true networks. Cosine distance values closest to 0 indicate the greatest possible similarity to the true networks. A) time-lagged bivariate cross-correlation, no comparisons significant; B) time-lagged partial cross-correlation, all comparisons significant except 100% vs. 90%, 90% vs 80%, 80% vs. 70%, 70% vs. 60%, 60% vs 50%, 50% vs 40%, 50% vs. 90%, 40% vs. 20%, 40% vs. 10%, 30% vs. 20%, and 20% vs. 10%; C) time-lagged multivariate Granger causality, all comparisons significant, excluding 100% vs. 90%, 100% vs. 80%, 90% vs. 80%, 80% vs. 70%, 70% vs. 60%, 60% vs. 10%, 50% vs. 40%, 50% vs. 30%, 50% vs. 20%, 50% vs. 10%, 40% vs. 30%, 30% vs. 20%, and 20% vs. 10%; D) time-lagged bivariate mutual information, no comparisons significant; E) time-lagged bivariate transfer entropy, no comparisons significant; F) time-lagged multivariate transfer entropy, all comparisons significant excluding 100% vs. 90%, 90% vs. 80%, 80% vs 70%, 70% vs. 60%, 60% vs. 50%, 50% vs. 40%, 50% vs. 30%, 50% vs. 10%, and 40% network coverage compared to all lower percentages of coverage; G) zero-lag bivariate cross-correlation, no comparisons significant; H) zero-lag partial cross-correlation, no comparisons significant; I) zero-lag bivariate mutual information, no comparisons significant; J) shuffled, no comparisons are significant except 90% vs. 10% coverage, 80% vs. 10%, 70% vs. 10%, 60% vs. 10%, and 40% vs. 10%. General patterns do not show large declines in reconstruction reliability for most metrics as the percentage of the network covered decreases. Exceptions include partial cross-correlation which shows significant worsening, and multivariate Granger causality, which demonstrates significant improvement.

With 90–100% coverage, multivariate transfer entropy and partial cross-correlation performed similarly (Supp. Fig. 4, p = 0.0046, p = 0.0011), with MVTE significantly outperforming all other metrics. As coverage declined to 80–60%, MVTE remained the most accurate method, performing significantly better than all other metrics (p < 0.002). At 50% network coverage, Granger causality matched the performance of MVTE, and at 10%, mutual information, Granger causality, and bivariate transfer entropy performed comparably to multivariate transfer entropy (p < 0.0001). For most metrics (excluding mutual information), reducing network coverage increased variance in reconstruction accuracy (Fig. 8, p < 0.0001). Similar trends were observed in shuffled conditions, with small networks exhibiting high variance. Zero-lag metrics consistently underperformed, yielding worse results than time-lagged metrics and performing comparably to shuffled networks (p < 0.0001).

### Runtime

The computational time required to calculate EC can impact a metric’s practical utility. Rapid computation is essential for the future of real-time applications, such as closed-loop neuromodulatory devices, whereas post-hoc analyses can tolerate slower methods. To assess efficiency, we measured the runtime for each metric, acknowledging that while absolute times may vary by computing environment, relative trends remain consistent.

Despite its strong performance, MVTE is computationally intensive. For 10-node networks, MVTE averages 10 minutes to compute, compared to 20 seconds for bivariate cross-correlation (28 times faster) and 10 seconds for mutual information (475 times faster) (Fig. 9). As network size increases, TE-based runtimes scale inefficiently compared to other metrics. In 65-node networks, MVTE takes over 40 hours to compute, while partial cross-correlation takes 15 minutes, and mutual information takes just 20 seconds—nearly 7,000 times faster than MVTE. While bivariate transfer entropy also has a high computational cost, it is slightly faster than multivariate TE.

**Fig. 9.**
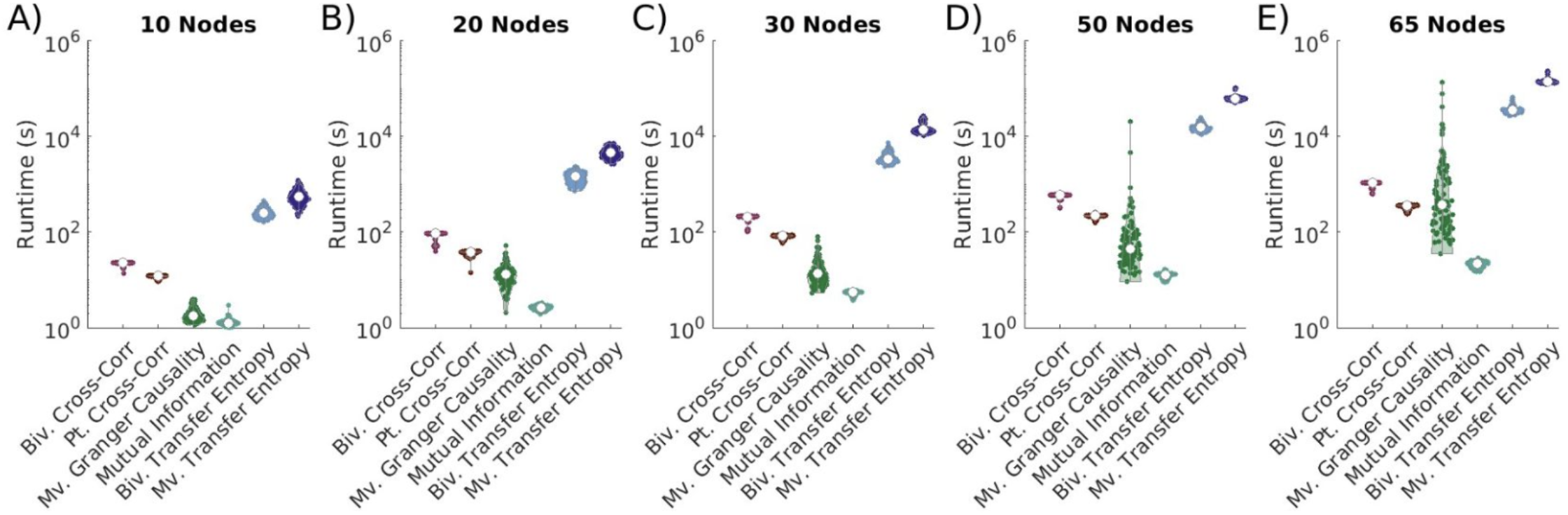
Comparative runtimes for EC metric calculation. Comparative runtimes on a Iog10-scale of time-lagged EC metrics for A) 10, B) 20, C) 30, D) 50, and E) 65 node networks. Results show that mutual information consistently has the fastest runtime among metrics used for this analysis, while bivariate and multivariate transfer entropy consistently have the longest runtimes.

Among non-TE metrics, mutual information is the most efficient, while bivariate cross-correlation is the slowest. Granger causality is often comparable to cross-correlation metrics.

## DISCUSSION

In these experiments, we examined several practical limitations encountered in intracranial electrophysiology recordings and their impact on the accuracy of network reconstructions. To accomplish this, we generated simulations of time series data with a vector autoregressive model using Erdős-Rényi connectivity. We then applied multiple EC metrics to infer network topology and evaluated their performance under different conditions.

In this study, our findings confirm that most EC metrics struggle to accurately reconstruct larger networks, with performance declining as the network size grows. The exception to this trend is partial cross-correlation, which improves with increasing network size. For small networks (<20 nodes), mutual information, Granger causality, and MVTE perform well, though MVTE’s computational demands remain a significant limitation. However, as network size grows, the accuracy of mutual information and Granger causality declines, while partial cross-correlation and MVTE maintain high performance, significantly outperforming other metrics in networks >20 nodes. Granger causality’s poor performance with larger networks is noteworthy, as it relies on an autoregressive framework, yet fails to scale effectively. Additionally, our results highlight that MVTE and partial metrics outperformed bivariate ones in networks of 30 nodes or more, emphasizing the importance of capturing multi-region interactions when analyzing large-scale brain networks.

### Effect of Network Size

Interactions across brain networks are crucial for cognitive function, often requiring high-density electrode arrays such as electrocorticography (ECoG) or stereoelectroencephalography (SEEG) to capture widespread activity. As the size of the networks increase, it is essential that EC metrics remain reliable in reconstructing connectivity. Our findings confirm that most EC metrics struggle to accurately reconstruct larger networks, with performance declining as the network size grows. The exception to this trend is partial cross-correlation, which improves with increasing network size. Partial cross correlation attempts to first account for the effect of other reference time-series before evaluating the relationship between the two nodes of interest. In data with a small number of nodes, these time processes are less likely to be normally distributed, and accounting for small skews in the data may induce the appearance of correlations.

For small networks (<20 nodes), mutual information, Granger causality, and MVTE perform well, though MVTE’s computational demands remain a significant limitation. However, as network size grows, the accuracy of mutual information and Granger causality declines, while partial cross-correlation and MVTE maintain high performance, significantly outperforming other metrics in networks >20 nodes. Granger causality’s poor performance with larger networks is noteworthy, as it relies on an autoregressive framework, yet fails to scale effectively. Additionally, our results highlight that MVTE and partial metrics outperformed bivariate ones in networks of 30 nodes or more, emphasizing the importance of capturing multi-region interactions when analyzing large-scale brain networks.

### Time series length

To assess how the length of time series impacts network reconstruction, we varied the number of time points available for analysis. While some network reconstructions, such as those from resting state data, may analyze extended recordings, many studies may investigate network connectivity changes over the course of a window within a behavioral trial. In these cases, selecting an appropriate EC metric is crucial for ensuring reliable network inference. As expected, our results demonstrate that reconstruction accuracy improves for all EC metrics as the number of time points increases. Among these, partial cross-correlation shows the greatest improvement with increasing time series length, suggesting its sensitivity to longer datasets. MVTE maintains consistent performance across all recording lengths, performing comparably to bivariate transfer entropy for shorter time series (500-1,000) points but excelling with longer recordings (10,000 points). These results highlight MVTE’s reliability when reconstructing networks, even with short recordings. One limitation of this analysis is its focus on time series length, excluding the potential influence of sampling frequency on reconstruction accuracy, where a basic assumption is that the sampling rate is sufficient to capture the dynamics of the system.

### Measurement Noise

Brain activity recorded through electrophysiology is often contaminated by multiple sources of noise in the environment, including movement artifacts, particularly in the operating room or hospital, where the presence of other medical equipment may increase the amount and number of sources of noise. While researchers take many precautions to reduce this noise, understanding how these metrics perform under noisy conditions is critical. Previous studies have emphasized the importance of characterizing and removing noise before network analysis (Bastos & Schoffelen, 2016; Pesaran et al., 2018), and others have established that standard techniques for noise removal, such as global signal regression (Saad et al., 2012) and filtering (Barnett & Seth, 2011) may introduce biases or distort network reconstructions.

Our analysis models signals with independent measurement noise sources for each electrode and assumes correlated noise had been removed. Several key findings emerged. First, and unsurprisingly, reducing noise levels improves the performance of all metrics, reinforcing the importance of both hardware-based noise reduction and analytical denoising techniques to enhance reconstruction validity. Second, MVTE remains robust across all noise levels and outperforms all other metrics. This aligns with findings from other experimental conditions about the superiority of MVTE for accurate network reconstructions. However, our analysis is limited by the assumption that noise sources are uncorrelated across nodes or that correlated noises cannot be removed, which may not be true for neural data with volume conduction.

### Network Coverage

In neural electrophysiology recordings, our electrodes often provide incomplete coverage of the networks of interest. Therefore, evaluating how EC metrics perform with missing data is essential for ensuring accurate network reconstructions. The impact of network coverage varies by metric. Bivariate cross-correlation, mutual information, and bivariate transfer entropy show minimal changes with declining coverage. MVTE exhibits slight declines but remains robust across all levels of coverage. In contrast, Granger causality and partial cross-correlation are more affected. Interestingly For Granger causality improves as network coverage declines with performance peaking at 50% (25 nodes). This aligns with our network size results, which show that Granger causality produces excellent network reconstructions for smaller networks of 10-20 nodes but poorly reconstructs larger networks of 30 nodes or larger. We see similar trends with declining network coverage in networks of other sizes, with Granger causality reconstructions improving until the networks are 20 nodes or smaller (not shown), suggesting that declining network coverage is likely an artifact of Granger causality’s inherent tendency to perform well for smaller networks given a fixed amount of time series data. Conversely, partial cross-correlation worsens significantly with reduced coverage, and it performs poorest when only 30% (15 nodes) of the network is included in reconstruction. This is consistent with network size results, which demonstrate that partial cross-correlation is unreliable for networks with fewer than 20 nodes. The pattern remains consistent across networks of varying sizes (not shown), suggesting the issue stems from node count rather than percent coverage.

Across all metrics, declining coverage also leads to greater variability in reconstruction accuracy, even for zero-lag metrics and shuffled data. This variability cannot be fully explained by network size changes, suggesting that missing nodes inherently increase reconstruction inconsistencies. One potential contributing factor not evaluated here may arise from the common input problem, where two nodes may share input from a single node. When not observed, multivariate methods may be unable to account for this connection, leading the EC metric to identify a spurious connection between two nodes that share an input. This is particularly relevant for intracranial networks, where studies show coverage is extremely limited, even in epilepsy patients with extensive montages (Anderson et al., 2021; Katz & Abel, 2019). These findings highlight the consistent reliability of EC metrics despite changes to the degree of network coverage, but the increased variability seen with declining coverage should be kept in mind when interpreting results from EC metrics.

### Zero-Lag Metrics

As expected, in networks with time-lagged relationships, zero-lag metrics (bivariate cross-correlation, partial cross-correlation, and bivariate mutual information) performed significantly worse than time-lagged metrics, but it is important to understand how poorly these metrics perform if time lags become nontrivial. In many cases, zero-lag metrics often fail to generate accurate networks, often performing no better than randomly shuffled network topologies (random). These findings highlight the necessity of using metrics that account for temporal delays in time-series data. In neural electrophysiology, where sampling rates often exceed the information transfer rates between brain regions, zero-lag metrics are generally unsuitable. For lower-frequency modalities like fMRI, researchers must consider whether expected changes span more than a single time sample when choosing between zero-lag versus lagged metrics. If network interactions evolve over timescales longer than a single sample, lagged metrics are likely essential for accurate reconstruction. These findings also emphasize the necessity of accounting for physiological communication lag between brain regions. Network connections may be missed if the analysis window is too short to capture the time required for neural signal propagation.

### Runtime

While MVTE outperforms other metrics in many contexts, its lengthy computational time limits its practical utility. For a 50-node network with 10 possible lags, TE averaged 17 hours, compared to just 12.8 seconds for mutual information, the fastest metric. As TE runtimes increase significantly with more lags, analyzing 20–50 node networks with a 2000 Hz sampling rate and physiological lags could take weeks, making TE impractical for real-time applications. For real-time or closed-loop applications, mutual information seems to offer the best balance between speed and accuracy. Additionally, partial cross-correlation is a viable alternative for larger networks, achieving comparable reconstructions while averaging 220.2 seconds for the same 50-node networks. Although partial cross-correlation is not ideal for real-time use, implementing it in lower level programming languages could enhance its feasibility for online applications.

## CONCLUSION

Effective connectivity metrics vary in their ability to reconstruct neural networks, making it crucial to understand their relative strengths and tradeoffs under different conditions. In this study, we systematically evaluated EC metrics on equal footing across key parameters, including network size, time series length, noise levels, and coverage, allowing for a direct comparison of their performance. Our results show that multivariate transfer entropy provides the most accurate reconstructions overall, but its high computational cost limits its practical use in real-time applications. In contrast, mutual information offers an efficient alternative, excelling in small to medium-sized networks with rapid computation times that make it suitable for online analyses. Partial cross-correlation performs well in large networks when sufficient time series data is available, providing reliable reconstructions with significantly shorter runtimes than MVTE. Importantly, zero-lag metrics consistently fail when network interactions occur across multiple time lags, underscoring the necessity of using time-lag sensitive EC methods in neural electrophysiology. By evaluating these metrics under matched conditions, this work clarifies the tradeoffs between accuracy, computational efficiency, and suitability for different datasets and approaches.

## ETHICS STATEMENT

This paper does not require ethical approval, since all data utilized for this paper are simulated.

## DATA AND CODE AVAILABILITY

All code for generating data and completing analysis for this paper is available at https://github.com/hermandarrowlab/ec-metrics-validation.

### External Software

Functional Connectivity Toolbox: https://sites.google.com/site/functionalconnectivitytoolbox/Multivariate Granger Causality Toolbox: https://www.mathworks.com/matlabcentral/fileexchange/78727-the-multivariate-granger-causality-mvgc-toolbox Fieldtrip Mutual Information: https://github.com/fieldtrip/fieldtrip/blob/master/connectivity/ft_connectivity_mutualinformation.m Information Dynamics Toolbox XL (IDTxl): https://github.com/pwollstadt/IDTxl

## AUTHOR CONTRIBUTIONS

*Conceptualization*: TIN, DPD

*Data Curation*: KD

*Formal Analysis*: KD, HF

*Funding Acquisition*: KD, DPD, ABH

*Investigation*: KD

*Methodology*: KD, DPD, TIN

*Project Administration*: DPD, TIN, ABH

*Resources*: DPD, TIN, ABH

*Software*: DPD, TIN, KD

*Supervision*: DPD, TIN, ABH

*Validation*: KD

*Visualization*: KD

*Writing – Original Draft Preparation*: KD, HF

*Writing – Review & Editing*: KD, HF, ABH, TIN, DPD

