## supplement for "Metric Validation for Detection of Delayed and Directed Coupling"

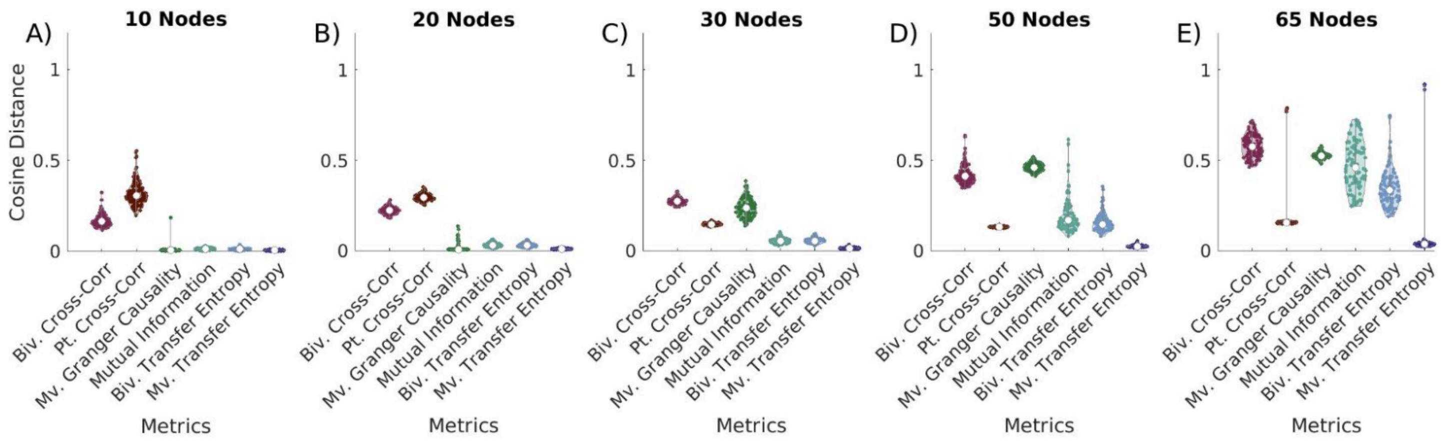


**Supplemental Fig. 1 - Effect of EC metric network on reconstruction in networks with increasing size.** Cosine distance values closest to 0 indicate the greatest possible similarity to true networks. Network similarity can be seen in networks of sizes A) 10 nodes, all comparisons significant excluding bivariate cross-correlation vs. partial cross-correlation, mutual information vs. multivariate Granger causality, mutual information vs. bivariate transfer entropy, multivariate Granger causality vs. bivariate transfer entropy, multivariate Granger causality vs. multivariate transfer entropy, and bivariate transfer entropy vs. multivariate transfer entropy; B) 20 nodes, all comparisons significant except bivariate cross-correlation vs. partial cross-correlation, mutual information vs. multivariate Granger causality, mutual information vs. bivariate transfer entropy, multivariate Granger causality vs. bivariate transfer entropy, multivariate Granger causality vs. multivariate transfer entropy; C) 30 nodes, all comparisons significant except BVXC vs. MVGC, PTXC vs. MI, PTXC vs MVGC, PTXC vs. BVTE, BVMI vs. BVTE, BVMI vs. MVTE, and BVTE vs. MVTE; D) 50 nodes, all comparisons significant except BVXC vs. MVGC, PTXC vs. MI, PTXC vs. BVTE, PTXC vs. MVTE, MI vs. BVTE; and E) 65 nodes, all comparisons significant except BVXC vs. BVMI, BVXC vs. MVGC, PTXC vs. BVTE, PTXC vs. MVTE, BVMI vs. MVGC, BVMI vs. BVTE, and MVGC vs. BVTE. As the number of nodes increases, there are changes to comparative metric performance. Multivariate transfer entropy is consistently a top performer across network sizes. In networks of 10-20 nodes, Granger causality performs comparably to multivariate transfer entropy. Bivariate transfer entropy also performs comparably to multivariate transfer entropy for networks of 10, 20 and 30 nodes, but worsens in comparison to multivariate transfer entropy in larger networks of 50 and 65 nodes. Partial cross-correlation improves with increasing network size, and for 50 and 65 node networks, this metric performs comparably to multivariate transfer entropy.

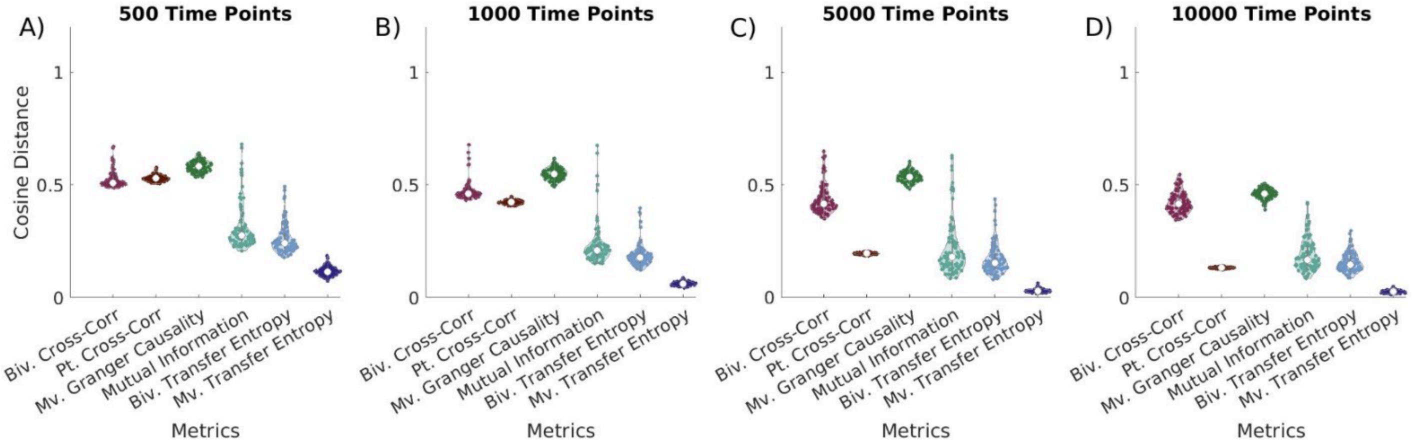


**Supplemental Fig. 2 - Effect of EC metric network reconstruction on ROC curves in networks with increasing numbers of time points** Cosine distance values closest to 0 indicate the greatest possible similarity to true networks. Network similarity can be seen in networks with A) 500 time points, all comparisons significant except BVXC vs. PTXC, BVXC vs. BVMI, BVXC vs. MVGC, PTXC vs. MVGC, BVMI vs. BVTE, and BVTE vs. MVTE; B) 1,000 time points, all comparisons significant except BVXC vs. PTXC, BVXC vs. MVGC, PTXC vs. BVMI, BVMI vs. BVTE, and BVTE vs. MVTE; C) 5,000 time points, all comparisons significant except BVXC vs. MVGC, PTXC vs. BVMI, PTXC vs. BVTE, BVMI vs. BVTE; D) 10,000 time points, all comparisons significant except BVXC vs. MVGC, PTXC vs. BVMI, PTXC vs. BVTE, PTXC vs. MVTE, and BVMI vs. BVTE. For 500 and 1,000 time point series, bivariate and multivariate transfer entropy perform comparably. At these lengths, multivariate transfer entropy significantly outperforms all other metrics, while bivariate transfer entropy also performs comparably to mutual information. At 5,000 points, multivariate transfer entropy outperforms all other metrics. At 10,000 time points, multivariate transfer entropy and partial cross-correlation perform comparably, and multivariate transfer entropy significantly outperforms all other metrics at this time series length. At 10,000 time points, partial cross correlation also performs comparably to Granger causality and mutual information.

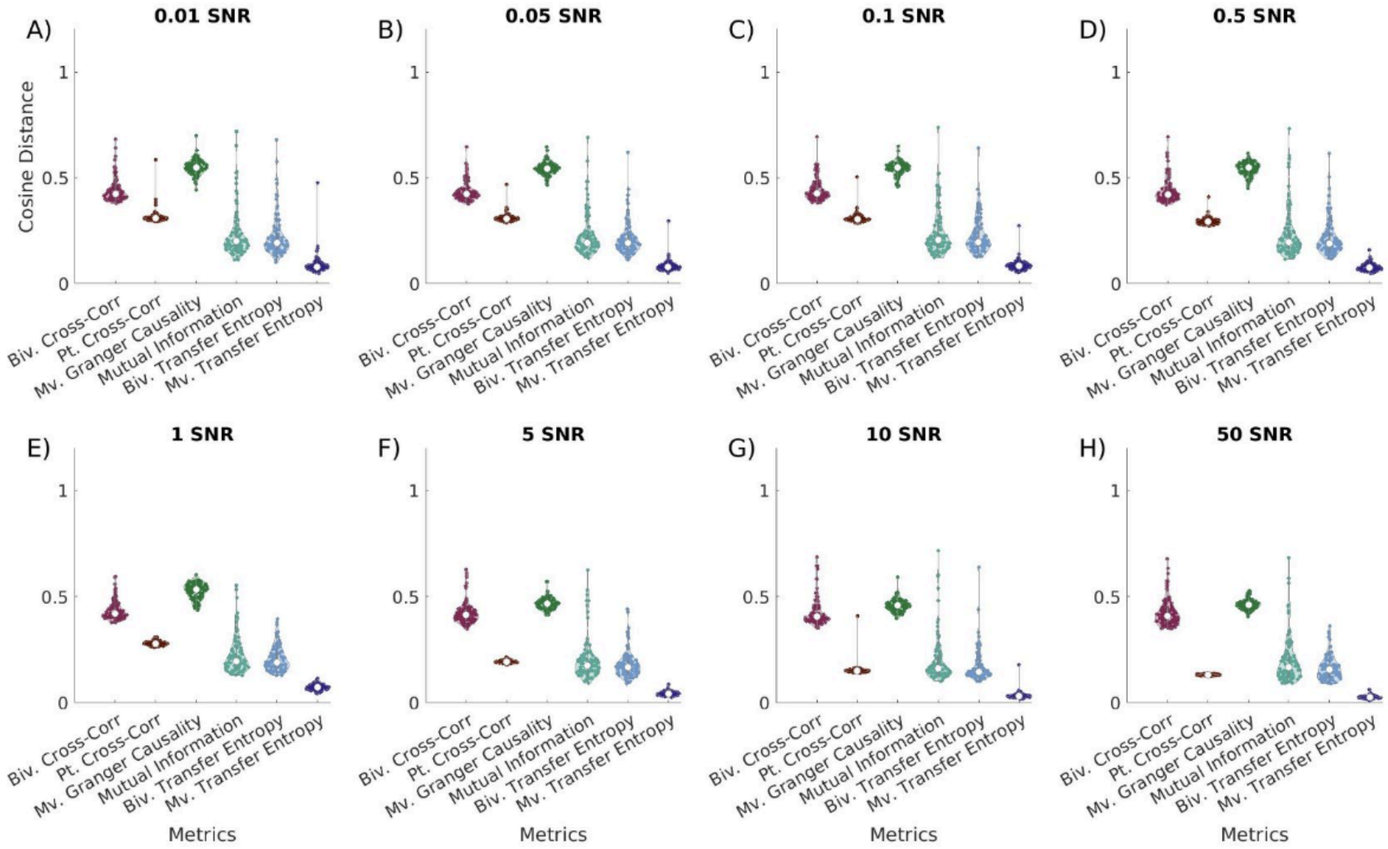


**Supplemental Fig. 3 - Effect of EC metric network on reconstruction in networks with increasing SNR.** Cosine distance values closest to 0 indicate the greatest possible similarity to the known networks. Network similarity can be seen in networks with A) SNR = 0.01, all comparisons significant except BVXC vs. PTXC, BVXC vs. MVGC, PTXC vs. BVMI, PTXC vs. BVTE, BVTE vs. MVTE; B) SNR = 0.05, all comparisons significant except BVXC vs. PTXC, BVCX vs. MVTE, PTXC vs. MVGC, BVMI vs. BVTE, BVMI vs. MVTE, and BVTE vs. MVTE; C) SNR = 0.1, all comparisons significant except BVXC vs. PTXC, PTXC vs. MVGC, BVMI vs. BVTE, BVMI vs. MVTE, BVTE vs. MVTE; D) SNR = 0.5, all comparisons significant except BVXC vs. PTXC, BVXC vs. MVGC, PTXC vs. BVMI, PTXC vs. BVTE, BVMI vs BVTE, BVMI vs. MVTE; E) SNR = 1, all comparisons significant except BVXC vs. MVGC, PTXC vs. BVMI, and PTXC vs. BVTE, BVTE vs. MVTE; F) SNR = 5, all comparisons significant except BVXC vs. MVGC, PTXC vs. BVMI, PTXC vs. BVTE, BVMI vs. BVTE; G) SNR = 10, all comparisons significant except BVXC vs. MVGC, PTXC vs. BVTE, and BVMI vs. BVTE; H) SNR = 50, all comparisons significant except BVXC vs. MVGC, PTXC vs. BVMI, PTXC vs. BVTE, PTXC vs. MVTE, BVMI vs. MVTE. For low SNR conditions, we see that multivariate transfer entropy performs comparably to bivariate transfer entropy, but significantly outperforms all other metrics. As noise increases, the performance of bivariate transfer entropy declines compared to multivariate transfer entropy. Multivariate transfer entropy outperforms bivariate transfer entropy when SNR = 0.1, 0.5, 5, 10, and 50. For all other noise conditions, multivariate transfer entropy significantly outperforms all other metrics. Additionally, bivariate transfer entropy, partial cross-correlation, and mutual information perform comparably for all noise levels.

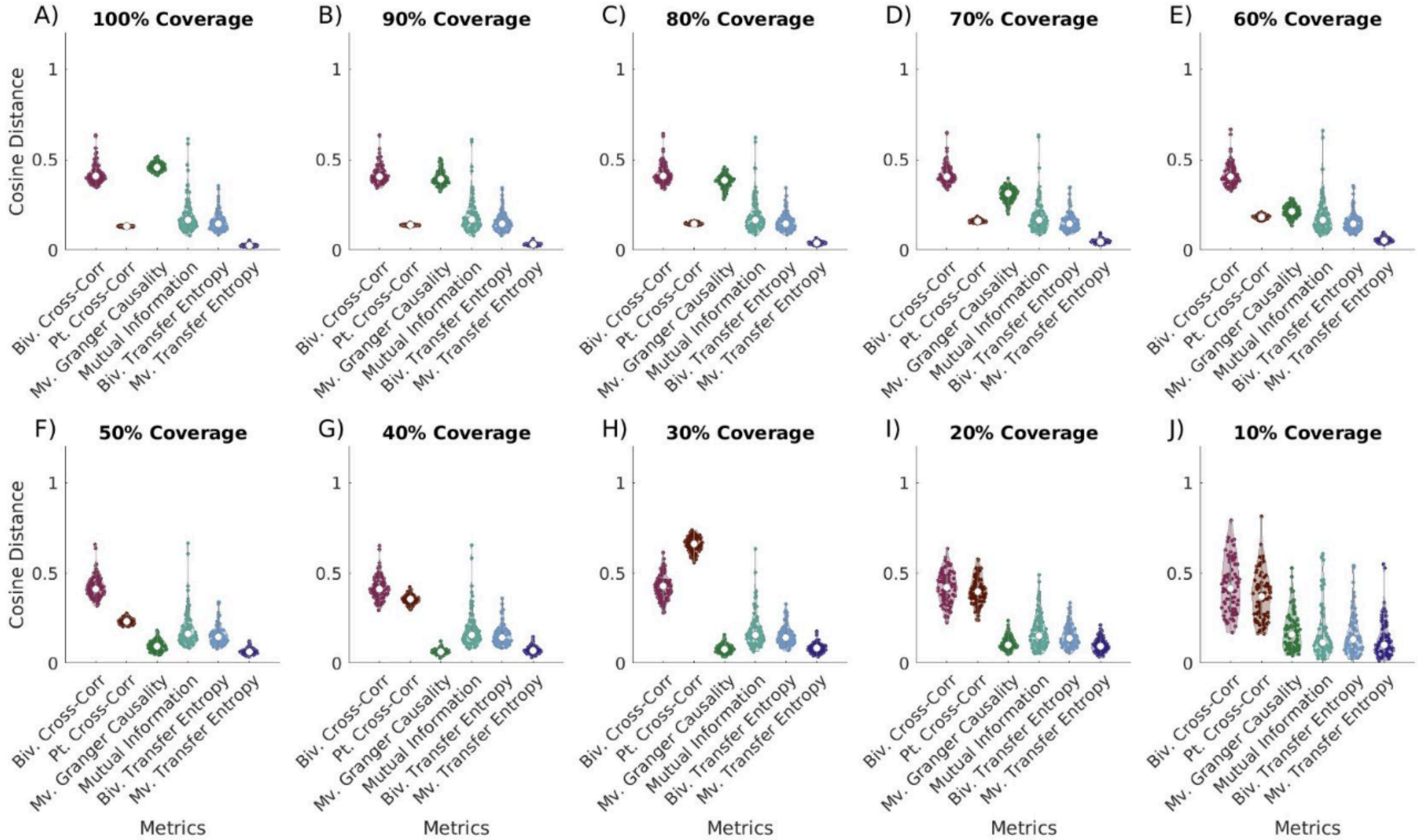


**Supplemental Fig. 4 - Effect of EC metric network reconstruction on networks with increasing white noise.** Cosine distance values closest to 0 indicate the greatest possible similarity to the true networks. Network similarity can be seen in networks with A) 100% network coverage, all comparisons significant except BVXC vs. MVGC, PTXC vs. BVMI, PTXC vs. BVTE, PTXC vs. MVTE, BVMI vs. BVTE; B) 90% Network Coverage, all comparisons significant except BVXC vs. MVGC, PTXC vs. BVMI, PTXC vs. BVTE, BVMI vs. BVTE; C) 80% network coverage, all comparisons significant except BVXC vs. MVGC, PTXC vs. BVMI, PTXC vs. BVTE, BVMI vs. BVTE; D) 70% network coverage, all comparisons significant except BVXC vs. MVGC, PTXC vs. BVMI, PTXC vs. BVTE, MVGC vs. BVMI, BVMI vs. BVTE; E) 60% network coverage, all comparisons significant except BVXC vs. MVGC, PTXC vs. MVGC, PTXC vs. BVMI, PTXC vs. BVMI, MVGC vs. BVMI, BVMI vs. BVTE; F) 50% network coverage, all comparisons significant except BVXC vs. PTXC, PTXC vs. BVMI, PTXC vs. BVTE, MVGC vs. BVTE, MVGC vs. MVTE, BVMI vs. BVTE; G) 40% network coverage, all comparisons significant except BVXC vs. PTXC, PTXC vs. BVMI, MVGC vs. BVTE, BVMI vs. BVTE; H) 30% network coverage, all comparisons significant except BVXC vs. PTXC, BVXC vs. BVMI, MVGC vs. BVTE, BVTE vs. MVTE; I) 20% network coverage, all comparisons significant except BVXC vs. PTXC, MVGC vs. BVMI, MVGC vs. BVTE, MVGC vs. MVTE, BVMI vs. BVTE, BVMI vs. MVTE, BVTE vs. MVTE; J) 10% network coverage, all comparisons significant except BVXC vs. PTXC, MVGC vs. BVMI, MVGC vs. BVTE, MVGC vs. MVTE, BVMI vs. BVTE, BVMI vs. MVTE, BVTE vs. MVTE. In conditions with 100% network coverage, partial cross-correlation and multivariate transfer entropy perform comparably. As network coverage declines, partial cross-correlations worsens, and multivariate granger causality improves. For lowest levels of network coverage (10-20%), non-cross-correlation metrics perform comparably.
